# Examining the genetic and phenotypic correlation between survival and fecundity in a wild bird

**DOI:** 10.1101/2025.05.14.653981

**Authors:** Lucy A. Winder, Joel Pick, Julia Schroeder, Mirre J.P. Simons, Terry Burke

## Abstract

There is intraspecific variation in reproduction and survival, which has been hypothesised to be the result of trade-offs in investment. However, we lack compelling evidence that trade- offs drive variation within species, perhaps because genetic trade-offs are masked by environmental variation. We used a long-term dataset of breeding records for a closed population of house sparrows (*Passer domesticus*) and its associated pedigree to separate the genetic and environmental covariance between fecundity and survival. We measured age-specific changes in reproductive outputs and estimated the heritability of both offspring production and survival in multivariate animal models. Our results show no evidence for survival costs to reproductive output. Nor did we find evidence that individuals’ fecundity and survival are correlated either the genetic or permanent environment level. Our findings show no evidence that variation in fecundity is driven by the offspring-production/survival trade-off. Instead, our results show that individuals are only somewhat consistent in their reproductive output, and suggest this is a possible mechanism why selection cannot act on individual quality.

## Background

Theory suggests that, as resources are limited, trade-offs in resource allocation will cause variation between individuals in their life histories, such as in fecundity, offspring quality and individual survival (Stearns 1976; van Noordwijk and de Jong 1986; Kirkwood and Rose 1991; LemaÎtre and Gaillard 2017). However, there is little evidence for the reduction in performance of traits that is expected as a result of increased allocation towards reproduction (Reznick 1985; Zera and Harshman 2001; Santos and Nakagawa 2012; Cohen et al. 2020; Chang et al. 2024; Winder et al. 2025). It has also been suggested that reproductive variation is driven by underlying differences in individual condition (van Noordwijk and de Jong 1986; Pettifor et al. 1988; Winder et al. 2025). As environmental effects can obscure genetic variation, by separating these directly in a quantitative genetics framework, it is possible to identify the otherwise elusive costs of reproduction.

If individuals in a given population have multiple fitness-related traits that are positively correlated - i.e., individual “quality” - and if these traits are heritable, we would expect “quality” to respond to selection. It is thus surprising that positive correlations between fitness related traits are observed as, the population should evolve towards higher performing traits. Heritability has been well documented for fecundity-related traits, showing that additive genetic variation persists (Kruuk et al. 2000; Teplitsky et al. 2009; Brommer et al. 2010; Schroeder et al. 2012b). By contrast, there has been mixed evidence of heritability of longevity (Merilï and Sheldon 2000; Kruuk et al. 2000; Schroeder et al. 2012b; Vedder et al. 2021a). It is therefore possible that environmental effects act in opposition to the genetic drivers of fitness-related traits, thereby keeping reproductive variation within a population. For example, if an individual has benefited from favourable early-life conditions, they may perform better without necessarily having a genetic predisposition to higher fitness and survival. There are several studies that provide evidence that an individual’s environment shapes its life-history trajectory and, thereby, leads to different phenotypic quality between individuals, for example: manipulation of early-life conditions (Alonso-Alvarez et al. 2006; Spagopoulou et al. 2020), lifetime manipulation of environmental conditions (Marasco et al. 2018), and observational evidence of varying environmental conditions (Pettorelli et al. 2001).

Why phenotypic correlations between fecundity and survival are not consistently observed across studies could be explained by terminal effects. Increases in reproductive output when individuals face impending mortality (for example, with severe disease), known as terminal investment, have been observed (for example, Bonneaud et al. 2004; Hanssen 2006).

However, reductions in reproductive output are also possible if an individual’s condition deteriorates prior to death (Coulson and Fairweather 2001; Rattiste 2004). These sudden reductions in reproductive performance in the final breeding attempt for individuals may be misinterpreted as increased senescence. Only in the last decade have long-term individual-based studies gained enough power to separate between-individual effects of ageing (Bouwhuis et al. 2009; Schroeder et al. 2012a; Hammers et al. 2012). However, a limiting factor for most of these studies remains that it is difficult to determine if individuals have died or have dispersed from the study area, meaning disentangling effects of survival remain largely uncorroborated. In this study, we quantify correlations between fecundity and survival in genetic and permanent environment space. We take advantage of a long-term individual-based study of a closed population of house sparrows (*Passer domesticus*) for which individuals in our study have accurate death records. We find that correlations observed in our dataset between fecundity and survival are driven by terminal effects. Therefore, we conclude that there is no evidence for genetic or environmental trade-offs.

## Materials and Methods

### Data collection

Data were collected on a population of house sparrows (*Passer domesticus*) on Lundy Island in the Bristol Channel (51º10’N, 4º40’W). The population has been closely monitored since 1996 but for the purpose of this analysis we excluded data from before 2001 to ensure that all records are from birds of known age and that the majority of the population is known (>99%). Individuals are ringed with a BTO metal ring, a unique combination of colour rings and are fitted with a passive-integrated transponder as either nestlings or fledglings, meaning that precise ages are known for all individuals. Nests were monitored throughout each breeding season, so that reproductive outputs for the majority of breeding attempts are known. As sparrows do not undertake long-distance flight, the population is considered to be closed to immigration and emigration (Schroeder et al. 2015). Therefore, we also have an accurate estimation of when an individual died. This information means we have complete life histories at an individual level. All individuals were blood sampled to form a genetic pedigree (all procedures were performed under UK Home Office licence). The genetic pedigree was assembled using 13 microsatellite loci to identify genetic parents (Dawson et al. 2012).

### Fitness measures

We used exclusively female fitness components in this analysis, measured as the number of broods produced in a year, the total number of eggs produced in a year and the number of nestlings per brood at five days post-hatching. If a bird was known to be alive in a year but had no breeding record, its fecundity was recorded as zero. However, if it was the last year in which a bird was known to be alive, fecundity was recorded as a missing value as we could not be certain of how many breeding attempts these individuals would have had (i.e., an individual could have died part way through the breeding season).

Individual survival records were defined by whether an individual survives to the following year. This was determined by either sightings at the nestbox during incubation or provisioning of chicks, or by capture during mist-netting sessions throughout the year (in both the summer breeding season and a winter monitoring period). Birds that are not sighted for two consecutive years are considered to have died after their last sighting. Previous work in our population has shown a resighting probability of 0.96 (Simons et al. 2015).

### Data analysis

All analyses were performed using R version 3.6.2 (R Development Core Team 2009).

Mortality risks as a function of offspring production were estimated using interval-based Cox proportional hazards models using the coxme package (Therneau 2020). These models were used to determine the relationship between offspring production and survival/mortality at its simplest level to illustrate how a perceived trade-off may be observed when other variables are not accounted for. The total number of eggs produced was modelled as a fixed effect and the focal year was a random effect in the model.

In order to estimate the genetic variance of fitness components, we analysed a) the total annual number of broods produced, b) the total annual number of eggs produced, and c) the total annual number of chicks produced alive at 5 days post hatching, using separate generalised linear animal models (MCMCglmm) with a gaussian distribution (Hadfield 2010). We included age and its squared term, maximum age and whether the breeding attempt was the individuals last (i.e., to test for a terminal effect) as covariates in our model (van de Pol and Verhulst 2006; van de Pol and Wright 2009). The permanent environmental effects and genetic effects were also included as random effects in the model. The genetic effects were modelled using a matrix for the additive genetic relationship between individuals, built from the pedigree. The focal year was also included as a random effect.

We also modelled survival probability using an approximation of the Cox Proportional Hazards (CoxPH) model (see Zhong et al. 2019). Survival was fitted with a Poisson distribution. Age was expressed as a categorical fixed effect, which is a function of how the CoxPH model is approximated. The results of this model were similar to modelling survival as a binomial variable and our conclusions would not have changed had we used this approach. As in the fecundity model, between-individual, genetic and year effects were modelled as random effects.

We ran bivariate models of reproductive effort and survival using MCMCglmm. This allowed us to explore whether there is a correlation between reproductive effort and survival and what drives this variation (i.e., a between-individual effect vs genetic effect). As we found similar effects for broods, eggs and chicks, in the univariate analysis of reproductive output we, again, used these three measures of reproductive effort a) the total annual number of broods produced, b) the total annual number of eggs produced, and c) the total annual number of chicks produced alive at 5 days post hatching, and all were modelled using a gaussian distribution. Survival was again estimated using an approximation of the Cox Proportional Hazards model, using a Poisson distribution. Age and its quadratic term were fixed effects for fecundity. We did not use max age as this is instead taken into account in the survival component of the model. We removed the categorical age effect for survival, as survival did not vary significantly between age groups and removing the effect did not change the conclusions drawn from our results. We modelled the permanent environmental effect, genetic effect and focal year effect as random effects in the model.

We repeated this analysis, but for the reproductive effort in the final year, if the bird was known to have an incomplete breeding season in the final year (i.e., it died part way through the season), we removed any reproductive output values, including these instead as missing information. A bird was considered to have an incomplete final breeding season if it was not sighted in August or later months of its last year. Sightings of individuals outside of the nest are largely derived from mist netting of adults and juveniles in early winter. In these models, we also included a terminal effect as a fixed effect.

The animal models for survival and reproduction were all run using MCMCglmm (version 2.34). For all models the burn-in period was 60,000, the chain length was 460,000 iterations and the thinning interval was 200. For the random effects, parameter expanded priors were used with a mean of 0 and variance (α.μ = 0, α.V = 1000 × I_2_). For residual variances, inverse wishart priors were used (V =1 and nu =0.002 for the univariate models, and V= diag(2) and nu = 1.002 for the bivariate models).

## Results

We found high reproduction is correlated with low mortality risk (Table 1). However, in our univariate model of reproductive output, we found within-individual changes with age follow the commonly found bell-shaped pattern with age (Bouwhuis et al. 2009; Hammers et al. 2012), with offspring production starting and accelerating from the first year of breeding but levelled off in mid life and decreasing through later life (Figure 1, Table 2). We did not find an effect of maximum age (Table 2), showing that individuals who live longer do not produce more offspring than those who do not live as long (Table 2). We found similar age-related fecundity patterns when using different fitness proxies (broods, eggs and chicks) (Table 2).

**Table 1:** Results of Cox hazards models for mortality given annual egg production. 960 observations from 495 individuals were used in this analysis.

|  | coef | exp(coef) | SE | z | p |
| --- | --- | --- | --- | --- | --- |
| Annual egg production | -0.032 | 0.968 | 0.013 | -2.46 | **0.014** |

**Table 2:** Model outputs for a univariate animal model of annual reproductive output (n = 996 observations from 507 individual females). The reproductive output measures used in this analysis are annual totals of broods, eggs and chicks at age 5 days post-hatching. Fixed effect values show the posterior mean (95% credible intervals). Values for random effects show the posterior mode of the proportion of variance explained by each variable (95% credible intervals). Bold text indicates statistical significance.

|  | Broods | Eggs | Chicks |
| --- | --- | --- | --- |
| Fixed: |  |  |  |
| (Intercept) | **1.87 (1.61-2.10)** | **6.57 (5.39-7.92)** | **2.99 (2.10-3.92)** |
| Age | **0.43 (0.26-0.59)** | **2.11 (1.39-2.82)** | **1.59 (1.11-2.14)** |
| Maximum age | 0.01 (-0.07-0.07) | -0.01 (-0.30-0.29) | 0.01 (-0.21-0.25) |
| Age^2 | **-0.05 (-0.07--0.03)** | **-0.23 -(0.33--0.13)** | **-0.17 (-0.24--0.10)** |
| Last year | **-0.36 (-0.50--0.18)** | **-1.48 (-2.23--0.82)** | **-0.65 (-1.20--0.13)** |
| Random: |  |  |  |
| Permanent environment | 0.14 (0.02-0.24) | 0.12 (0.00-0.23) | 0.13 (0.06-0.27) |
| Genetic ($h^2$) | 0.00 (0.00-0.13) | 0.05 (0.00-0.20) | 0.00 (0.00-0.13) |
| Focal year | 0.03 (0.01-0.11) | 0.07 (0.03-0.18) | 0.04 (0.02-0.13) |
| Residual | 0.75 (0.65-0.84) | 0.67 (0.57-0.77) | 0.70 (0.62-0.81) |

**Figure 1:**
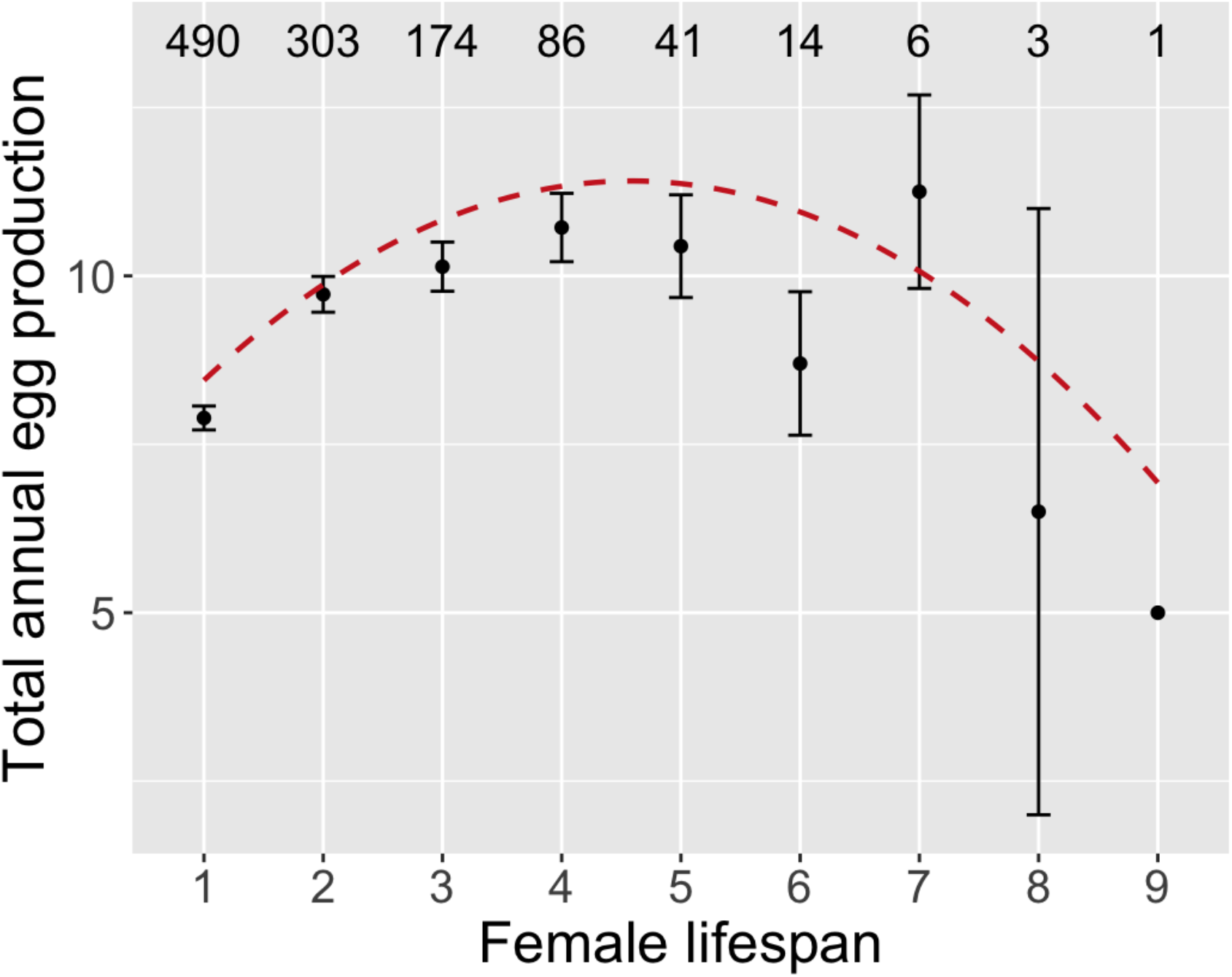
Total annual offspring production (number of eggs produced) given the lifespan of the female parent. Points are the population mean offspring production for the given lifespan. Whiskers are the standard error from the mean. The curves represent the within-individual change in offspring production as females age (from model outputs in Table 2). Sample sizes are given above the plot for each lifespan.

### Heritability of fecundity and survival

We only found minimal heritability of annual egg production (*h*^*2*^ = 0.05, Table 2), but on no other fecundity trait. We also found a permanent environmental effect on annual brood, egg and chick production (proportion of variance = 0.14, 0.12 and 0.13 respectively). We found negligible heritability of survival (*h*^*2*^ = 0.01 (95% CI: 0.00-0.01), Table 3). Survival did not differ significantly between age groups (Table 3). Focal year explained the most variance in survival probability.

**Table 3:** Model outputs for a univariate model of survival probability. Survival was estimated for each age using an approximation of the Cox proportional hazards model. 507 females for this analysis.

|  | Posterior mean | 95% CI - lower | 95% CI - upper | pMCMC | Proportion of variance |
| --- | --- | --- | --- | --- | --- |
| Fixed: |  |  |  |  |  |
| Age 1 | -0.74 | -1.09 | -0.46 | <0.001 |  |
| Age 2 | -0.63 | -0.96 | -0.29 | <0.001 |  |
| Age 3 | -0.74 | -1.12 | -0.39 | <0.001 |  |
| Age 4 | -0.78 | -1.21 | -0.39 | <0.001 |  |
| Age 5 | -1.00 | -1.55 | -0.38 | <0.001 |  |
| Age 6 | -1.11 | -2.16 | -0.05 | 0.02 |  |
| Age 7 | -0.61 | -1.86 | 0.70 | 0.33 |  |
| Age 8 | -1.43 | -4.00 | 1.00 | 0.31 |  |
| Age 9 | -15.00 | -32.92 | 1.67 | 0.04 |  |
| Random: |  |  |  |  |  |
| Between-individual | 0.00 | 0.00 | 0.01 |  | 0.01 |
| Genetic | 0.00 | 0.00 | 0.01 |  | 0.01 |
| Focal year | 0.39 | 0.09 | 0.78 |  | 0.97 |
| Residual | 0.00 | 0.00 | 0.01 |  | 0.01 |

### Fecundity-survival correlation

Using multivariate models, we found a positive, but not statistically-significant, permanent environment correlation between annual reproductive output and survival, where individuals that produced more offspring had a higher survival probability (Figure 2, Supplementary Information S2, S4 and S6). However, we found no evidence of a genetic correlation between reproduction and survival in any fecundity measure (Figure 2, Supplementary Information S2, S4 and S6). We did find a significant positive correlation between annual brood production and survival at the focal year level (Figure 2, Supplementary Information S2), though the trend was positive but non-significant for annual egg production and annual number of chicks (Figure 2, Supplementary Information S4 and S6 respectively). At the residual level, we found a significant positive correlation between reproduction and survival for all reproductive measures (Figure 2, Supplementary Information S2, S4 and S6). For all reproductive measures, we found moderate permanent environment variation in fecundity and small genetic variation in fecundity (Figure 2, Supplementary Information S2, S4 and S6). Variation in survival was almost exclusively explained by focal year (Figure 2, Supplementary Information S2, S4 & S6). Similarly to the univariate model, we found that as age increased, fecundity output (for all reproductive measures) also increased, but decreased later in age, indicating senescence in fecundity (Supplementary Information S1, S3 and S5).

**Figure 2:**
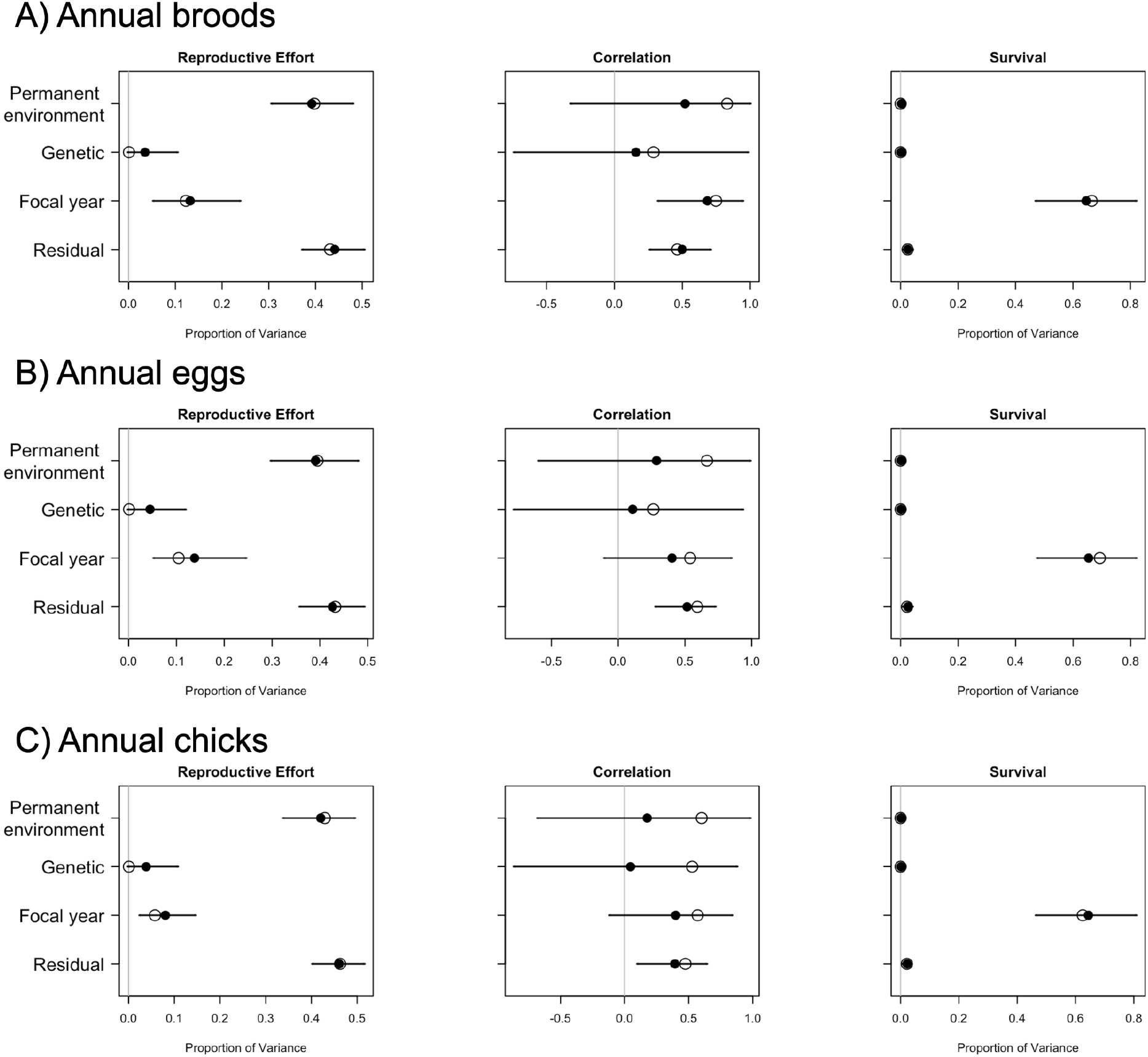
The correlation between fecundity (A) reported as total annual brood production, B) reported as total annual egg production and C) reported as total annual chicks at age 5 days post-hatching) and survival, and the proportion of variance of both traits. The order of effects (from top down) is the same for fecundity and survival as shown for the correlation. Filled circles are the mean and open circles are the mode variance of the posterior MCMC samples. Whiskers are the 95% credible intervals. Note that a positive correlation indicates that higher brood production predicted higher survival probability.

When we included a terminal effect (whether it was an individual’s last breeding year) and incomplete final breeding records (accounted for by removing the fecundity data for individuals who had an incomplete breeding record in their final year) this reduced the permanent environment correlation between fecundity and survival to zero for all reproductive measures (Figure 3, Supplementary Information S2, S4 and S6). The genetic correlation also remained at zero. We found no significant correlation between any reproductive measure and survival at the focal year and residual levels (Figure 3, Supplementary Information S2, S4 and S6). However, for the number of chicks, the focal year correlation was positive, though non-significant (Figure 3, Supplementary Information S6). The terminal effects detected in all the reproductive measures were positive, meaning individuals in their year before their deaths showed increased reproductive output (Supplementary Information S1, S3 and S5). The difference in sign in the terminal effect between the univariate and multivariate analyses is driven by the removal of incomplete breeding records in the final year of breeding for the multivariate analysis.

**Figure 3:**
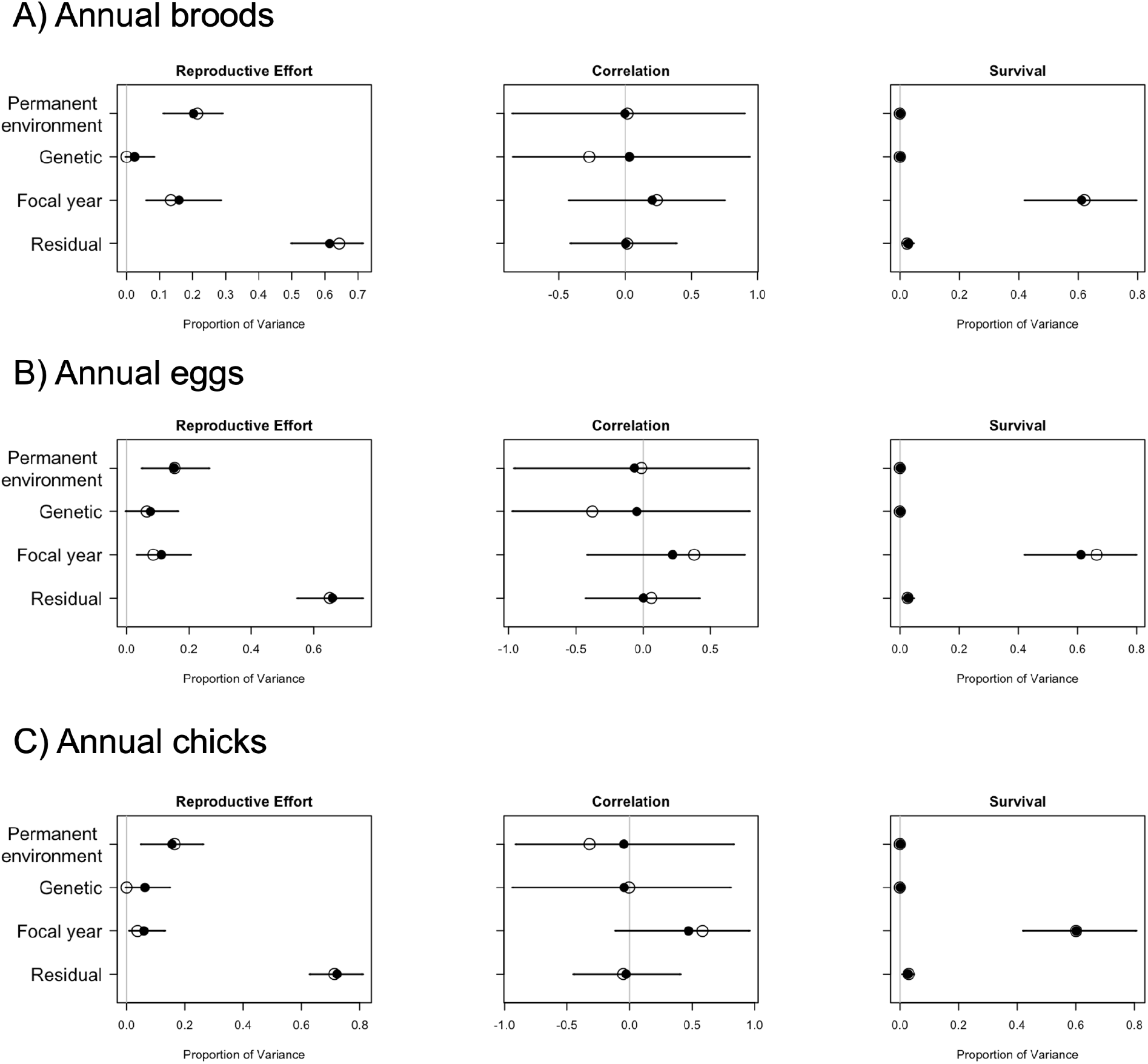
Model outputs for bivariate models of fecundity (A) reported as total annual brood production, B) reported as total annual egg production and C) reported as total annual chicks at age 5 days post-hatching) and survival. The dataset used has fecundity records removed in the least year an individual is alive if the individual has an incomplete final breeding season. A terminal effect has also been included as a fixed effect associated with fecundity. The outputs show the correlation between fecundity and survival, and the proportion of variance of both traits. The order of effects (from top down) is the same for fecundity and survival as shown for the correlation. Filled circles are the mean and open circles are the mode variance of the posterior MCMC samples. Whiskers are the 95% credible intervals. Note that a negative correlation indicates that lower brood production had a lower survival probability.

Any evidence of a correlation between survival and reproduction, therefore, appears driven by terminal or age effects between individuals. Individuals still show a repeatability of 0.23 for brood production, 0.23 for egg production and 0.22 for chick production, and variance in both reproduction and survival is driven in part by year (Figure 3, Supplementary Information S2, S4 and S6). Annual fecundity was shown to have a small heritability of 0.03 (0.00-0.08) for brood production, 0.07 (0.00-0.16) for egg production and 0.06 (0.00-0.15) for chick production (using the bivariate analyses including selective disappearance and terminal effect). However, the permanent environment effects explained more variation in fecundity than the genetic effects (0.20, 95% CI: 0.12-0.30 for brood production, 0.16, 95% CI: 0.05- 0.27 for egg production and 0.16, 95% CI: 0.05-0.26 for chick production). However, as the residual variation in fecundity explained more variation than either permanent environment and genetic variation (0.62, 95% CI: 0.50-0.72 for number of broods, 0.66, 95% CI: 0.56- 0.76 for the number of eggs and 0.72, 95% CI: 0.63-0.81), individual quality can be considered to not always be consistent.

## Discussion

Our results demonstrate the importance of accounting for age-specific changes in fecundity. We detected a negative trend between fecundity and mortality risk but found no correlation between fecundity and survival in the bivariate analysis at the permanent environment or genetic level. This difference is probably due to the modelling of age-associated changes in fecundity. As birds aged, the number of offspring they produced increased, whilst at older ages, there is evidence of late-life senescence. However, we found that when between- and within-individual age-related changes in reproductive output are considered, it becomes apparent that it is individuals that live longer that produce more offspring annually. Lundy sparrows, therefore, at the population level, show a commonly found pattern of senescence, with increases in reproductive output in early life followed by a steady decline in late life (e.g., Bouwhuis et al. 2009).

A key finding of this work is that a positive (though non-significant) correlation between survival and annual reproductive output reduces to zero when both terminal investment and individuals with an incomplete final breeding season are accounted for (note this also explains the negative terminal effect observed in the univariate fecundity model). This raises an important issue for many wild studies - observed correlations between life-history traits should be treated with caution, as the correlation is potentially driven by unexplained variation. When accounting for individuals that do have a complete breeding season in their final year, we found evidence for terminal investment where reproductive output increases in the last year an individual is alive. The reason for this is possibly one of resource allocation. An individual at the end of its life will not benefit from allocating resources to self-maintenance, when this will not ensure the individual will survive to the following breeding season (Bonneaud et al. 2004; Hanssen 2006; Froy et al. 2013; Simons et al. 2016). Instead, a better strategy would be to invest heavily in reproduction to maximise the genetic contribution to the future population. Without accounting for selective disappearance, we would have concluded that individuals have a terminal decline in reproductive effort in their final year for which we were able to determine through sighting efforts both within and outside of the breeding season. Furthermore, without accounting for either selective disappearance or a terminal effect, we would have concluded there is evidence that variation in fecundity is driven by differences in individual quality.

We have found that the costs of reproduction are minimal, counter to the currently favoured hypothesis that life-history trade-offs drive variation in fecundity (Stearns 1976, though see Cohen et al. 2020). In none of the models fitted was there evidence for a trade-off. It is possible that our data did not have the power to detect this effect, which could explain the large credible intervals observed. However, even if this were the case, it is unlikely that the selection pressure would be large enough to produce any phenotypic change in the population. A recent study by Chang et al. (2024) also highlighted that genetic trade-offs are often absent. Importantly, our study fails to detect a trade-off between reproduction and survival in a wild dataset using modern quantitative genetics methods. Our results, therefore, further emphasise that there is limited evidence for trade-offs operating in the wild (Winder et al. 2025).

One explanation for the lack of apparent costs of reproduction is that individuals differ in their ability to acquire resources, leading to positive correlations (van Noordwijk and de Jong 1986). Our results show no (or negligible) among-individual correlation between fecundity and survival. This shows that an individual is not consistent in their quality across traits. For example, older individuals may be advantaged with experience when environmental conditions are unfavourable (Lunn et al. 1994), whereas in other years senescence would cause them to have lower reproductive output than younger birds in the population (Hammers et al. 2012). Consistency would be expected, however, when quality is viewed as a collection of traits that make an individual predisposed to perform above the population average at a given time, be that in fecundity, survival and/or access to desired resources. In that framework, quality represents the exact opposite of a trade-off, with traits being positively as opposed to negatively correlated. It is instead likely that there are complex interactions between individual condition and the environment, meaning life-history traits are stochastic.

Another (but not mutually exclusive) explanation for our results showing no costs of reproduction is offspring variability. As our results showed no genetic correlation between fecundity and survival, no heritability of survival and small heritability but a significant permanent environment effect of fecundity, we can conclude that offspring are variable in their fitness. Why this is remains unclear. One possibility is that trade-offs occur within the nest through limits on the amount of parental care that can be given (e.g., provisioning). Another possibility is that offspring vary in their life-history strategy (Wolf and McNamara 2012). Creating variable offspring within a brood has the potential to increase the likelihood of the parent having a successful breeding attempt (i.e., having at least one offspring recruit) if environmental conditions are unfavourable. There is growing evidence that offspring within the same brood follow alternative life-history trajectories dictated by their specific developmental conditions (Slagsvold et al. 1984; Groothuis et al. 2005; Muller and Groothuis 2013; Drummond and Rodríguez 2013; Vedder et al. 2021b).

## Conclusions

Using a long-term individual-based dataset, we found no evidence for a survival cost to reproduction. The alternative hypothesis, individual quality, was not supported as we did not detect a positive correlation between fecundity and survival. Only a small proportion of the variation in fecundity was attributable to individuals being consistent in their output.

Therefore, individuals are not consistent in their quality relative to the population and thus traits that contribute to an individual reproductive performance at one time point are selected against at another time point. We consider environmental variation and changing selection pressures the most likely reasons why variation in reproduction and possibly reproductive strategies are maintained. We suggest that this force is stronger and possibly more relevant to understanding ecology and evolution than trade-offs.

## Author contributions

LAW: Conceptualization, data curation, formal analysis, investigation, methodology, visualisation, writing - original draft. JP: Formal analysis, methodology, visualisation, writing - review & editing. JS: Conceptualization, Writing - review & editing. MJPS: Conceptualization, riting - review & editing, supervision. TB: Funding acquisition, writing - review & editing, supervision.

## Supplementary Information

**S1:** Model output for the fixed effects in bivariate models of fecundity, measured as annual number of broods produced, and survival. Survival was modelled as an approximation of the Cox proportional hazards model. 507 females were used for these models. The associated random effects for these models can be found in S2.

| Model |  | Posterior mean | 95% CI - lower | 95% CI - upper | pMCMC |
| --- | --- | --- | --- | --- | --- |
| A.<br>Base model | Intercept | 1.93 | 1.72 | 2.14 | <0.001 |
|  | Age | 0.32 | 0.22 | 0.41 | <0.001 |
|  | Age<sup>2</sup> | -0.03 | -0.05 | -0.02 | <0.001 |
| B.<br>Selective disappearance data:<br>Base model + terminal effect | Intercept | 1.43 | 1.04 | 1.76 | <0.001 |
|  | Age | 0.47 | 0.30 | 0.65 | <0.001 |
|  | Age<sup>2</sup> | -0.05 | -0.08 | -0.02 | 0.002 |
|  | Terminal effect | 0.25 | 0.05 | 0.45 | 0.008 |

S2: The variance components predicting fecundity (annual number of broods produced) and survival (n = 507 females) and the correlation between the two traits estimated in a bivariate model. Values are the posterior means with credible intervals in parentheses. The results for the main effects of these models can be found in S1.

**S2:** The variance components predicting fecundity (annual number of broods produced) and survival (n = 507 females) and the correlation between the two traits estimated in a bivariate model. Values are the posterior means with credible intervals in parentheses. The results for the main effects of these models can be found in S1.

|  |  | Proportion of Variance |  |  |
| --- | --- | --- | --- | --- |
| Model |  | Total annual brood production | Survival | Correlation |
| A.<br>Base model | Between-individual | 0.39<br>(0.30-0.48) | 0.00<br>(0.00-0.01) | 0.54<br>(-0.27-0.99) |
|  | Genetic | 0.03<br>(0.00-0.10) | 0.00<br>(0.00-0.01) | 0.15<br>(-0.73-0.98) |
|  | Year | 0.13<br>(0.05-0.23) | 0.65<br>(0.48-0.82) | 0.68<br>(0.33-0.93) |
|  | Residual | 0.44<br>(0.37-0.50) | 0.02<br>(0.01-0.04) | 0.49<br>(0.26-0.69) |
| B.<br>Selective disappearance data:<br>Base model + terminal effect | Between-individual | 0.20<br>(0.12-0.30) | 0.00<br>(0.00-0.00) | 0.00<br>(-0.89-0.89) |
|  | Genetic | 0.03<br>(0.00-0.08) | 0.00<br>(0.00-0.00) | 0.03<br>(-0.84-0.93) |
|  | Year | 0.16<br>(0.07-0.28) | 0.62<br>(0.45-0.81) | 0.20<br>(-0.42-0.72) |
|  | Residual | 0.62<br>(0.50-0.72) | 0.03<br>(0.01-0.04) | 0.00<br>(-0.36-0.40) |

S3: Model output for the fixed effects in bivariate models of fecundity, measured as total annual egg production, and survival (n = 507 females). Survival was modelled as an approximation of the Cox proportional hazards model. The results of the associated random effects can be found in S4.

**S3:**
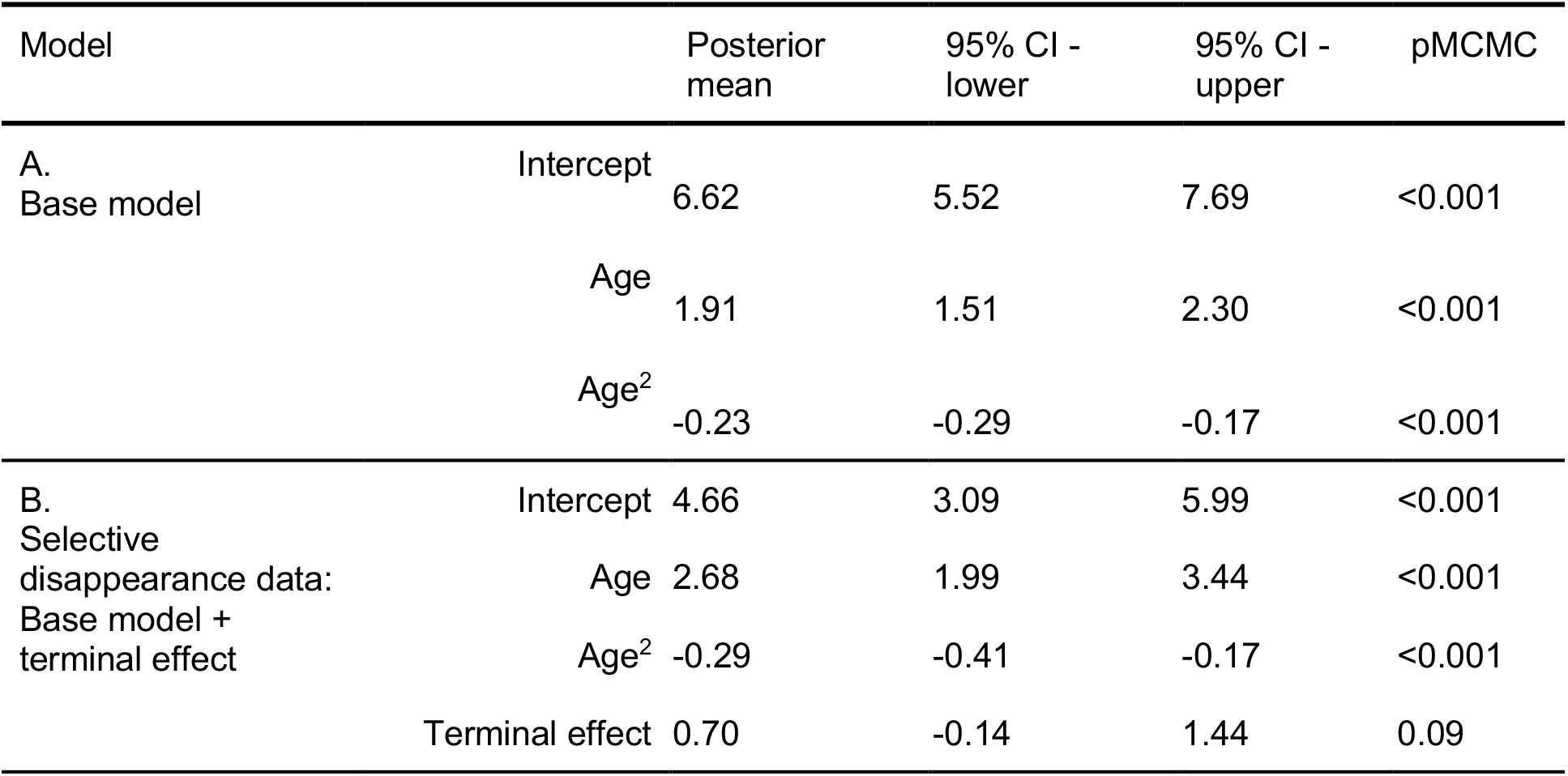
Model output for the fixed effects in bivariate models of fecundity, measured as total annual egg production, and survival (n = 507 females). Survival was modelled as an approximation of the Cox proportional hazards model. The results of the associated random effects can be found in S4.

**S4:** Variance components from bivariate models predicting fecundity (total annual egg production) and survival, and the correlation between the two traits (n = 507 females). Values are the posterior means with credible intervals in parentheses. The main effects from these models can be found in S3.

| Model |  | Proportion of Variance |  |  |
| --- | --- | --- | --- | --- |
|  |  | Total annual egg production | Survival | Correlation |
| A.<br>Base model | Between-individual | 0.39<br>(0.30-0.48) | 0.00<br>(0.00-0.01) | 0.29<br>(-0.59-0.99) |
|  | Genetic | 0.04<br>(0.00-0.12) | 0.00<br>(0.00-0.01) | 0.11<br>(-0.78-0.93) |
|  | Year | 0.14<br>(0.05-0.24) | 0.65<br>(0.48-0.82) | 0.40<br>(-0.10-0.84) |
|  | Residual | 0.43<br>(0.36-0.49) | 0.02<br>(0.01-0.04) | 0.52<br>(0.28-0.72) |
| B.<br>Selective disappearance data:<br>Base model + terminal effect | Between-individual | 0.16<br>(0.05-0.27) | 0.00<br>(0.00-0.01) | -0.08<br>(-0.91-0.80) |
|  | Genetic | 0.07<br>(0.00-0.16) | 0.00<br>(0.00-0.00) | -0.03<br>(-0.92-0.82) |
|  | Year | 0.11<br>(0.04-0.20) | 0.61<br>(0.43-0.78) | 0.22<br>(-0.34-0.76) |
|  | Residual | 0.66<br>(0.56-0.76) | 0.03<br>(0.01-0.05) | -0.01<br>(-0.41-0.38) |

S5: Model output for the fixed effects in bivariate models of fecundity, measured as total annual number of chicks at age 5 days post-hatching, and survival (n = 507 females). Survival was modelled as an approximation of the Cox proportional hazards model. The results of the associated random effects can be found in S6.

**S5:**
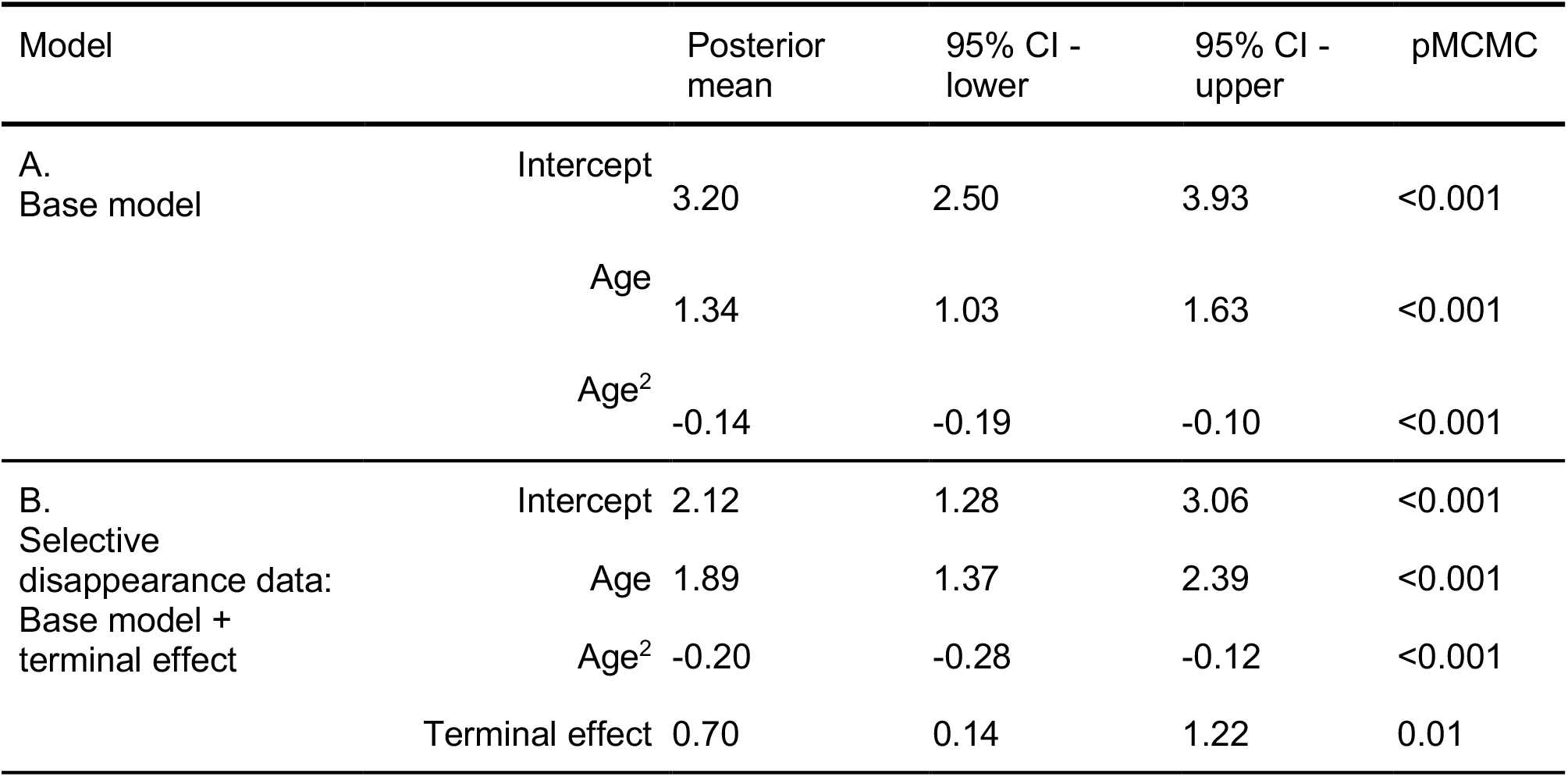
Model output for the fixed effects in bivariate models of fecundity, measured as total annual number of chicks at age 5 days post-hatching, and survival (n = 507 females). Survival was modelled as an approximation of the Cox proportional hazards model. The results of the associated random effects can be found in S6.

| Model |  | Posterior mean | 95% CI - lower | 95% CI - upper | pMCMC |
| --- | --- | --- | --- | --- | --- |
| A.<br>Base model | Intercept | 3.20 | 2.50 | 3.93 | <0.001 |
|  | Age | 1.34 | 1.03 | 1.63 | <0.001 |
|  | Age<sup>2</sup> | -0.14 | -0.19 | -0.10 | <0.001 |
| B.<br>Selective disappearance data:<br>Base model + terminal effect | Intercept | 2.12 | 1.28 | 3.06 | <0.001 |
|  | Age | 1.89 | 1.37 | 2.39 | <0.001 |
|  | Age<sup>2</sup> | -0.20 | -0.28 | -0.12 | <0.001 |
|  | Terminal effect | 0.70 | 0.14 | 1.22 | 0.01 |

S6: Variance components from bivariate models predicting fecundity (total chicks at age 5) and survival, and the correlation between the two traits (n = 507 females). Values are the posterior means with credible intervals in parentheses. The main effects from these models can be found in S5.

**S6:** Variance components from bivariate models predicting fecundity (total chicks at age 5) and survival, and the correlation between the two traits (n = 507 females). Values are the posterior means with credible intervals in parentheses. The main effects from these models can be found in S5.

|  |  | Proportion of Variance |  |  |
| --- | --- | --- | --- | --- |
| Model |  | Total annual chicks age 5 | Survival | Correlation |
| A.<br>Base model | Between-individual | 0.42<br>(0.39-0.49) | 0.00<br>(0.00-0.01) | 0.17<br>(-0.68-0.98) |
|  | Genetic | 0.04<br>(0.00-0.11) | 0.00<br>(0.00-0.00) | 0.05<br>(-0.86-0.87) |
|  | Year | 0.08<br>(0.03-0.15) | 0.64<br>(0.47-0.81) | 0.40<br>(-0.11-0.84) |
|  | Residual | 0.46<br>(0.40-0.51) | 0.02<br>(0.01-0.04) | 0.39<br>(0.10-0.64) |
| B.<br>Selective disappearance data:<br>Base model + terminal effect | Between-individual | 0.16<br>(0.05-0.26) | 0.00<br>(0.00-0.01) | -0.05<br>(-0.90-0.83) |
|  | Genetic | 0.06<br>(0.00-0.15) | 0.00<br>(0.00-0.01) | -0.04<br>(-0.93-0.80) |
|  | Year | 0.06<br>(0.01-0.13) | 0.60<br>(0.42-0.80) | 0.47<br>(-0.11-0.95) |
|  | Residual | 0.72<br>(0.63-0.81) | 0.03<br>(0.01-0.05) | -0.03<br>(-0.44-0.40) |

